# Spatiotemporal transcriptional dynamics of the cycling mouse oviduct

**DOI:** 10.1101/2021.01.15.426867

**Authors:** Elle C. Roberson, Anna M. Battenhouse, Riddhiman K. Garge, Ngan Kim Tran, Edward M. Marcotte, John B. Wallingford

## Abstract

Female fertility in mammals requires iterative remodeling of the entire adult female reproductive tract across the menstrual/estrous cycle. However, while transcriptome dynamics across the estrous cycle have been reported in human and bovine models, no global analysis of gene expression across the estrous cycle has yet been reported for the mouse. Here, we examined the cellular composition and global transcriptional dynamics of the mouse oviduct along the anteroposterior axis and across the estrous cycle. We observed robust patterns of differential gene expression along the anteroposterior axis, but we found surprisingly few changes in gene expression across the estrous cycle. Notable gene expression differences along the anteroposterior axis included a surprising enrichment for genes related to embryonic development, such as Hox and Wnt genes. The relatively stable transcriptional dynamics across the estrous cycle differ markedly from other mammals, leading us to speculate that this is an evolutionarily derived state that may reflect the extremely rapid five-day mouse estrous cycle. This dataset fills a critical gap by providing an important genomic resource for a highly tractable genetic model of mammalian female reproduction.

## Introduction

The adult mammalian female reproductive organs – ovary, oviduct, uterus, cervix, and vagina – hold an interesting position in mammalian physiology because they constantly and repeatedly engage in a complex remodeling process more commonly associated with development. With every menstrual or estrous cycle, fluctuations in ovarian steroid hormone secretion drive remodeling of these tissues, characterized by changes in proliferation, apoptosis, epithelial morphology, and fluid secretion (Brenner and West, 1975). However, while decades of research have revealed how these hormones fluctuate across the menstrual/estrous cycle in multiple mammals, we understand far less about the mechanisms by which these hormones drive cyclic tissue morphogenesis.

The oviducts are most well-known as the conduit between the ovary and uterus, but it is critical to note they also function as the site of fertilization and pre-implantation embryonic development (Coy et al., 2012; Coy and Yanagimachi, 2015; Stewart and Behringer, 2012). Interestingly, the oviducts display robust patterning along the anteroposterior axis, as the two major epithelial cell types – multiciliated cells (MCCs) and secretory cells – are differentially enriched (Agduhr, 1927; Barton et al., 2020; Stewart and Behringer, 2012). At the anterior oviduct close to the ovary, MCCs are highly enriched and responsible for capturing the ovulated oocyte(s), while at the posterior close to the uterus, MCCs are very sparse and act as sperm reservoirs (Suarez, 2016; Talbot et al., 1999). The reduced proportion of MCCs in the posterior oviduct reflects the significant increase in the proportion of secretory cells, which are critical to pre-implantation development (Coy and Yanagimachi, 2015). This anteroposterior pattern of the mouse oviduct epithelium is known to be established early in postnatal life and requires signaling from the underlying mesenchyme (Yamanouchi et al., 2010). How this pattern may change across the estrous cycle is not known.

In many species, the cellular basis of morphogenesis of oviduct MCCs across the estrous cycle was described *via* scanning electron microscopy studies decades ago (Brenner, 1969; Shirley and Reeder, 1996; Verhage et al., 1973). In Rhesus monkeys, for example, anterior oviduct epithelial cells are cuboidal at the beginning of the cycle, but then grow in height and develop cilia at their apical surfaces as estrogen increases (Brenner, 1969). As estrogen decreases and progesterone increases, the MCCs regress and de-ciliate (Brenner, 1969).

In mice, however, such cellular changes have not been thoroughly investigated, although it was shown that the wet weight of oviducts, as well as RNA and protein concentration, increases in the first half of the estrous cycle, and then subsequently decreases during the second half (Bronson and Hamilton, 1971; Yamanouchi et al., 2010). More recently, there is evidence that ciliary beat frequency changes in response to estrogen and progesterone (Bylander et al., 2010; Shi et al., 2011). Perhaps surprisingly, while steroid hormone signaling in the oviduct is crucial for female fertility (Herrera et al., 2020; Winuthayanon et al., 2015), we still lack a comprehensive view of how cycling steroids impact cellular morphogenesis in the oviduct, especially in the mouse.

A major hurdle to filling this knowledge gap is our very limited understanding of the transcriptional dynamics that underlie cyclical morphogenesis in this tissue. Transcriptomic approaches in human, swine, and bovine oviducts have investigated the impact of estrogen and progesterone, and some studies have also explored anteroposterior patterning in humans and bovine oviducts (Bauersachs et al., 2004; Cerny et al., 2015; Hess et al., 2013; Kim et al., 2018; Sowamber et al., 2020). However, the majority of these studies have focused on either the transcriptional response to fertilization and early embryo development or transcriptional signatures associated with progression of high grade serous ovarian carcinoma in the oviduct. The absence of a dynamic mouse oviduct transcriptome across the normal estrous cycle represents a critical gap in our knowledge, especially given its potential utility as a model for understanding mammalian fertility.

Here, we examined both the cellular changes and global transcriptional dynamics along the length of the oviduct and across the estrous cycle. First, we provide quantitative data on the density of MCCs along the anteroposterior axis of the oviduct and we show that MCCs do not remodel across the estrous cycle in mice. In addition, we present 3’ RNA-seq (Lohman et al., 2016) data for the anterior (approximate infundibulum) and posterior (approximate isthmus) oviduct at each stage of the estrous cycle. While transcript abundances vary strongly along the anteroposterior axis, our analyses suggest the estrous cycle has a surprisingly limited impact on transcription. Our data complement previous studies of transcriptional dynamics in the oviduct of other mammals and provide an important new resource for genetic studies of oviduct function in the mouse.

## Materials and Methods

### Mice

6-8-week-old Swiss Webster female mice were obtained from Charles River and allowed to acclimate from travel for 1 week. Mice were housed in individually ventilated cages in a pathogen-free facility with continuous food and water, with a controlled light cycle (light from 7am-7pm). 7-9-week-old females were estrous cycle staged using standard vaginal cytology (Ajayi and Akhigbe, 2020). Mice were humanely euthanized with extended CO_2_ exposure followed by cervical dislocation, and female reproductive tracts were dissected. All animal experiments were approved by the University of Texas at Austin Institutional Animal Care and Use Committee.

### Tissue processing & immunofluorescence

Dissected oviducts were carefully linearized (Fig. 1A) by teasing apart the supercoils, gently affixing to a strip of index card to keep the tissue straight, then fixing in 4% paraformaldehyde for either 4-6hr at room temperature (RT) or overnight at 4°C. Linearized fixed oviducts were washed in PBS, and then incubated in 30% sucrose at 4°C overnight. Following a brief incubation in NEG-50 Frozen Section Medium (ThermoFisher), oviducts were embedded in NEG-50 using an ethanol/dry ice bath. 12μm frozen longitudinal sections were cut on a cryostat (Leica) and dried overnight at RT. Frozen sections were stored at −20°C.

**Figure 1.**
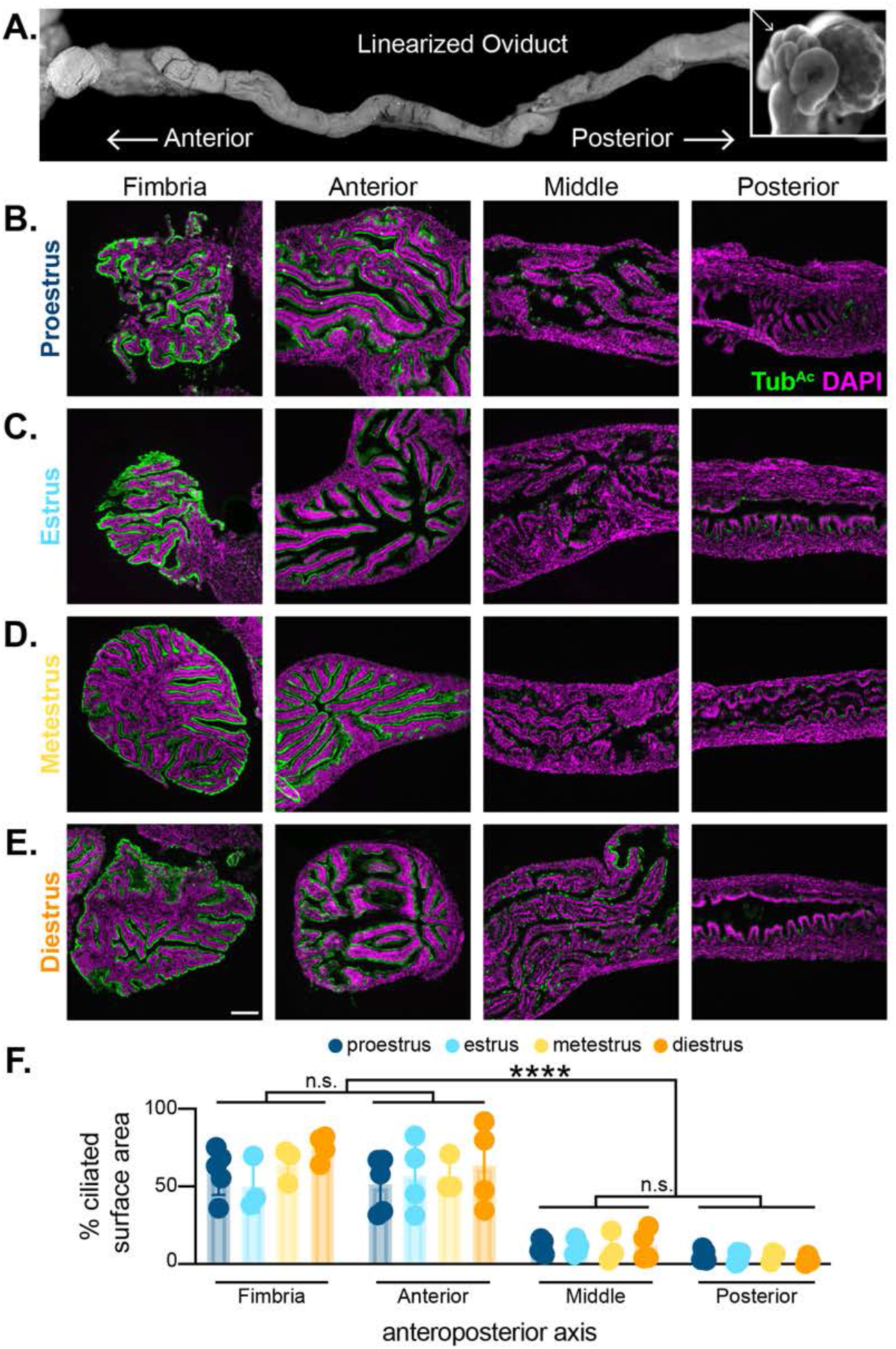
The mouse oviduct displays anteroposterior patterning that does not change across the estrous cycle. A) A linearized mouse oviduct, where the anterior is close to the ovary and the posterior is close to the uterus. The insert shows a non-linearized supercoiled mouse oviduct (insert, arrow). Oviducts were collected at B) proestrus, C) estrus, D) metestrus, and E) diestrus, linearized, and imaged for nuclei (DAPI, magenta) and cilia (Tub^Ac^, green) along the AP axis (fimbria, anterior, middle, and posterior). Scale bar = 100μm. F) Quantitation of the percent ciliated surface area along the oviduct lumen in each of the four estrous stages. Some error bars are too small to be seen. * = p< 0.05.

Tissue sections were washed in PBS + 0.1% Tween20 (PBST) three times to remove NEG-50. Tissues were blocked for at least 30min at RT with 5% normal donkey serum + PBS (block buffer). Primary antibodies for cilia (mouse anti-acetylated tubulin, 1:1000 dilution, Sigma, cat# 6-11B-1) were diluted in block buffer and incubated on slides for 2hr at RT or overnight at 4°C. After washing three times with PBST, slides were incubated with Alexa-Fluor coupled secondary antibodies (goat anti-mouse 647, 1:1000 dilution, ThermoFisher) and DAPI (1:1000 dilution, ThermoFisher) for at least 30min at RT. After washing, slides were mounted with Prolong Gold (ThermoFisher), and allowed to cure at RT in the dark overnight. Oviduct sections were imaged on either a Zeiss LSM700 point scanning confocal microscope or a CSU-W1 spinning disk Nikon confocal.

### Tissue sectioning and quantitation

Two to four sections spaced at least 100μm apart were analyzed for cilia at four locations along the anteroposterior axis of the oviduct. Images from the the fimbria, anterior third (approximate infundibulum), middle third (approximate ampulla), and posterior third (approximate isthmus) were acquired and quantified. We quantified ciliary area as previously published (Roberson et al., 2020). Briefly, we used FIJI image processing to trace the lumen based on DAPI staining to approximate the luminal length. Similarly, the ciliated surface of the lumen was traced and measured based on acetylated tubulin staining. From these measurements, the % ciliated surface area was calculated for each image, and we plotted using GraphPad Prism 8.

### RNA isolation and cDNA synthesis

For 3’ TagSeq (see below), dissected oviduct samples were collected in duplicate for each of the four stages across both anterior and posterior regions. For qPCR, samples were collected in triplicate for each estrous cycle stage across anterior, middle, and posterior thirds of the mouse oviduct. Following storage in RNA*Later* Storage Solution (Sigma, cat#: R0901) at −20°C, oviduct tissue was manually disrupted and the lysate was spun through a QIAshredder column (Qiagen, cat#: 79656) to fully homogenize. A Qiagen RNeasy mini kit (Qiagen, cat#: 74106) was used to harvest RNA for RNAseq, and the Qiagen RNeasy micro kit (Qiagen, cat#: 74004) was used to harvest RNA for qPCR. Total RNA was then either provided to the Genomic Sequencing and Analysis Facility at the University of Texas at Austin for 3’ TagSeq, or cDNA was synthesized using the iScript Reverse Transcription SuperMix (BioRad, cat#: 1708841) for qPCR.

### qPCR

Most primers were designed from a database for mouse and human qPCR primers incorporated into the UCSC genome browser (Zeisel et al., 2013). Primers for *Foxj1* and *Msx2* were designed using Primer3Plus software. See Supplemental Table S-1 for primer details. We confirmed specificity of primers by ensuring that they BLAST to no more than 1 site in the genome. There were four outliers to this BLAST assessment: all four ribosome primer sets blasted to more than 1 site in the genome, likely because there are numerous ribosomal pseudogenes scattered throughout the genome (Sisu et al., 2020). In addition, we only used primers whose melting curve displayed a single peak.

Primer sets were diluted from a stock (100μm in TE buffer) to 1μM in distilled deionized H_2_O. 2μL of each primer set (in technical duplicates) was allowed to dry in the bottom of a well in a MicroAmp Fast Optical 96 Well Reaction Plate (Thermofisher, cat#: 43-469-06). 10μL of a master mix of cDNA (250pg/well), Applied Biosystems SYBR Select Master Mix (Thermofisher, cat#: 44-729-18), and distilled deionized water was added to each well, the plates were sealed with MicroAmp Optical Adhesive Film (Thermofisher, cat#: 43-119-71) and allowed to incubate at RT in the dark for at least 15min to rehydrate primer. Plates were run on a ViiA-7 Real-Time PCR system (ThermoFisher), and CT values were auto-determined by the ViiA-7 software. The standard 2^−ΔΔCt^ method was then used to determine fold change based on the geomean of three ‘housekeeping’ genes (*Hprt*, *Dolk*, and *Sra1*) (Schmittgen and Livak, 2008).

### TagSeq

Tissue samples were collected in duplicate for each of the four estrous stages across both anterior and posterior regions of the mice oviduct, accounting for 16 samples in total. Library preparation and sequencing for TagSeq (Lohman et al., 2016; Meyer et al., 2011), a form of 3’ RNA sequencing, were performed by the Genomic Sequencing and Analysis Facility (GSAF) at The University of Texas at Austin. Total RNA was isolated from each sample by addition of Trizol (Thermo Fisher) and the sample was transferred to a Phasemaker tube (Thermo Fisher). Total RNA was extracted following the protocol supplied by the manufacturer and further cleaned up using a RNeasy MinElute Cleanup Kit (Qiagen). RNA integrity number (RIN) was measured using an Agilent Bioanalyzer and 100 ng of RNA was used for the TagSeq protocol. The fragmentation/RT mix was prepared and added to each RNA sample, then heated to 95°C for 2.5 minutes on a Thermocycler and immediately put on ice for 2 minutes. After cooling and addition of the template switching oligo and SmartScribe RT, the fragmented RNA reaction was incubated at 42°C for 1hr, 65°C for 15 min. Next an AmPure bead clean-up was completed for the cDNA before it was amplified to incorporate the Illumina sequencing primer site, followed by another cleanup. The remaining portions of the Illumina adapter (the i5 and i7 indices) were then added through an additional 4 cycles of PCR. Final libraries were quantified with PicoGreen then pooled equally for size selection using the Blue Pippin from 355-550 bp. Resulting libraries were sequenced using an Illumina HiSeq 2500 instrument (50-nt single reads). Full sample dataset details are provided in Supplemental Table S-2.

### Sequence data pre-processing

Fastq datasets were initially processed to collapse duplicates based on TagSeq molecular barcodes (Matz). Sequencing data quality, both before and after TagSeq pre-processing, was evaluated using the FastQC tool (v0.11.9) (Andrews) and reports were aggregated with the MultiQC program (v1.0) (Ewels et al., 2016).

### TagSeq data analysis

Single-end pseudo-alignment was performed against the mouse transcriptome (GENCODE M23 transcript sequences (Frankish et al., 2019)) using kallisto (v0.45.0) (Bray et al., 2016) with options -l 200 -s 50 --single-overhang -bias. Downstream analysis of transcript abundance data was performed in R (v3.4.4) following protocols outlined in Bioconductor (Gentleman et al., 2004). The tximport package (v1.6.0) (Soneson et al., 2015) was first used to roll up transcript-level counts into gene-level counts provided to the DESeq2 package (v1.18.1) (Love et al., 2014). Before further analysis, count data matrices were filtered to remove genes with fewer than 1 read across all included samples. A number of models were analyzed to explore the oviduct location/estrous stage relationship: Posterior versus Anterior locations, first providing Early and Late data separately (n=8 each), then again providing all datasets (n=16); and Late versus Early, first providing Anterior and Posterior location data separately (n=8 each), then again providing all datasets (n=16). Differentially expressed gene results reported are those with maximum adjusted p-value 0.05 and log2 fold change greater than 1.0 or less than −1.0.

Gene ontology (GO) analysis was performed using topGO R package (v2.34.0) (Alexa, 2020) with GO database org.Mm.eg.db (v3.7.0). The topGO classic algorithm and Fisher’s exact test were used in count data mode. Input genes had maximum adjusted p-values of 0.10. Separate analyses were performed for up-regulated (log2 fold change 0.5 or higher) and down-regulated (log2 fold change −0.5 or lower) genes. The background gene universe consisted of observed genes used in the DESeq2 analysis.

Full DESeq2 and topGO results are provided in the supplementary zip file for GEO accession number GSE164718.

## Results and Discussion

### Quantification of multiciliated cell density in the cycling mouse oviduct

Oviducts in most mammals consist of two key epithelial cell types – secretory and multiciliated cells (MCCs) – and these are unevenly distributed across the anteroposterior axis. MCCs are enriched anteriorly and secretory cells, posteriorly (Agduhr, 1927; Yamanouchi et al., 2010). While dynamic morphology changes across the estrous cycle have been described in the oviducts of many mammals (Brenner, 1969; Ferenczy et al., 1972; Novak and Everett, 1928; Shirley and Reeder, 1996; Verhage et al., 1973), the issue has not been investigated thoroughly in the mouse. To stringently assess cellular remodeling of the anteroposterior axis of the mouse oviduct across the estrous cycle, we linearized the normally supercoiled oviduct by carefully teasing apart each coil along the length of the organ (Fig. 1A), and generated longitudinal sections of linearized tissue. We then performed immunostaining for acetylated tubulin (Tub^Ac^) to label MCC cilia, and DAPI to label nuclei (Fig. 1B-E).

By examining fimbria, anterior (approximate infundibulum), middle (approximate ampulla), and posterior (approximate isthmus) regions of the oviduct, we clearly observed the anterior bias in MCC density (Fig. 1B-E). We quantified this pattern by calculating the percentage of cilia that line the oviduct lumen, based on Tub^Ac^ staining as described previously (Roberson et al., 2020) (Fig. 1F).

By performing parallel analyses on linearized oviducts from each stage of the estrous cycle, we found that this pattern displayed no temporal changes, with the anterior oviduct being significantly enriched for MCCs at all stages (Fig. 1F). The absence of significant remodeling of MCCs across the estrous cycle in the mouse oviduct presents a marked contrast to other mammals (Brenner, 1969; Shirley and Reeder, 1996; Verhage et al., 1973). We hypothesize that this discrepancy may relate to the very short (4 to 5 day) mouse estrous cycle (Ajayi and Akhigbe, 2020) and time required to eliminate and re-establish ciliated cells.

### Determining spatiotemporal transcriptome dynamics in the mouse oviduct

Patterns of secretion, and bulk RNA and protein concentrations are known to change during the mouse estrous cycle (Bronson and Hamilton, 1971). To ask if such changes have a transcriptional basis, we performed 3’ TagSeq, a high-throughput RNAseq profiling method that captures and sequences the 3’ ends of mRNA transcripts, enabling efficient estimation of relative gene expression (Lohman et al., 2016; Meyer et al., 2011). We performed two biological replicates of 3’ TagSeq on isolated anterior (highly ciliated) and posterior (minimally ciliated) oviducts at each stage of the estrous cycle (Fig. 2A). Principal component analysis (PCA) revealed that the majority of expression differences occur along the AP axis (72% of variance), while the estrous cycle accounts for much less variance (13%) (Fig. 2B).

**Figure 2.**
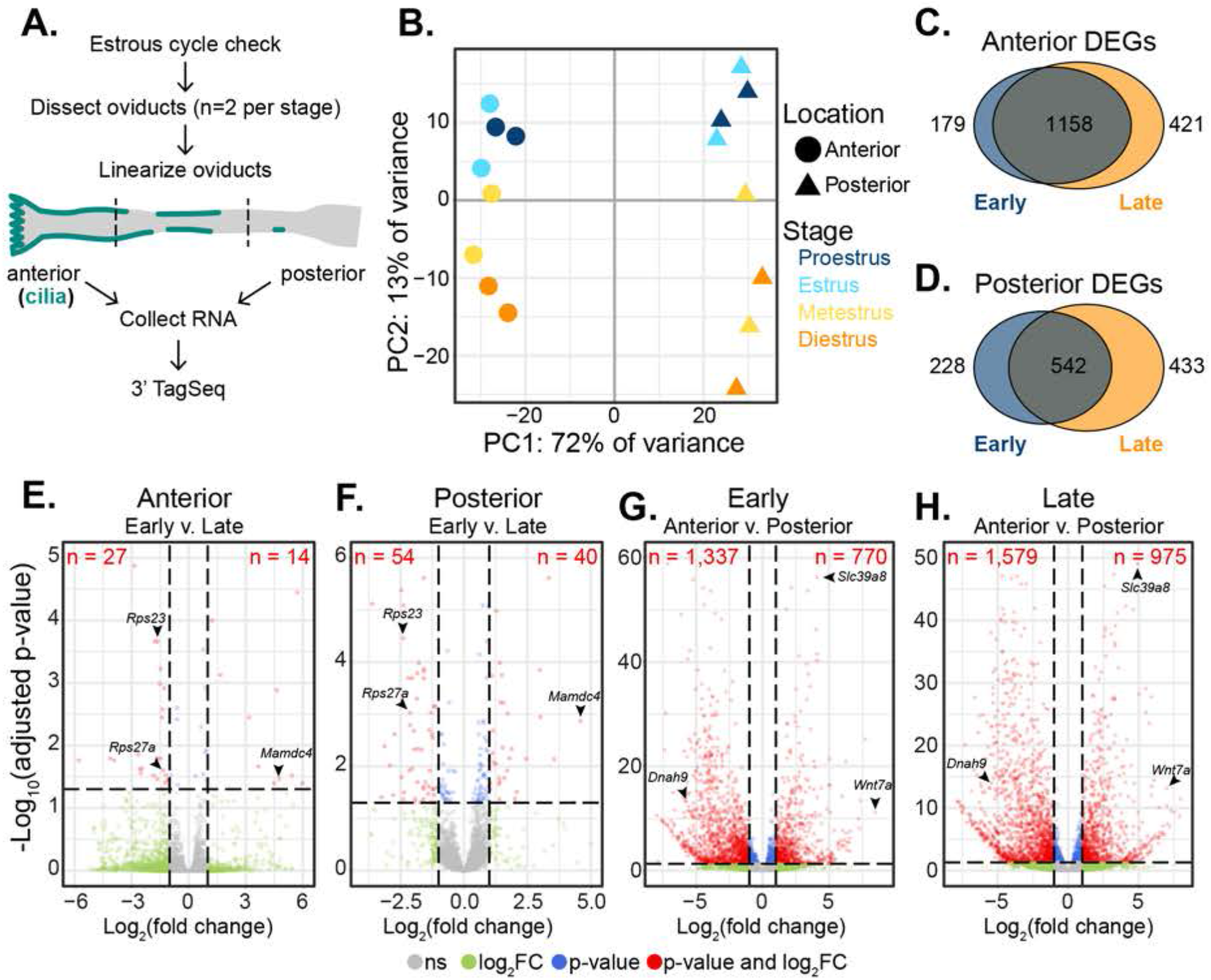
RNAseq of the oviduct shows major differences along the anteroposterior axis and minor change across the estrous cycle. A) Oviducts were dissected and linearized from each stage of the estrous cycle (n=2 each stage). RNA from the anterior and posterior thirds of each oviduct was collected and submitted for 3’ TagSeq. B) Principal Component Analysis (PCA) of the sixteen 3’TagSeq datasets. AP location (displayed as symbols) accounts for most of the variance while estrous cycle stage (displayed as colors) accounts for a lesser part with good separation between early (Proestrus, Estrus) and late (Metestrus, Diestrus) phases. C) Overlap between the anterior DEGs at early and late estrous cycle phases. D) Overlap between the posterior DEGs at early and late phases of the estrous cycle. Volcano plots of oviduct E) anterior and F) posterior compare early and late DEGs. Volcano plots of estrous cycle G) early and H) late phases cycle show anterior and posterior DEGs. Red points are genes significant at both adjusted P-value (<= 0.05) and effect size (log2 fold change <= −1 or >= 1).

The lack of strong variation across the estrous cycle was striking, so we considered that our data might be underpowered to detect statistically significant trends across time (n=2 per estrous cycle stage). We therefore repeated our analyses, collapsing the estrous cycle stages into ‘Early’ (proestrus and estrus) and ‘Late’ (metestrus and diestrus). This approach improved differentiation between the major cycle phases; the majority of differentially expressed genes (DEGs) were still found along the AP axis, not across the estrous cycle. For example, of the 1,758 DEGs enriched in the anterior oviduct, over 65% of them (1,158) were shared between Early and Late phases (Fig. 2C). Similarly, of the 1,203 DEGs enriched in the posterior, 45% (542) were shared between Early and Late (Fig 2D).

A more granular view of these data using volcano plots illustrates this finding. In the anterior oviduct, we identified only 27 early and 14 late DEGs (Fig. 2E). Similarly, modest enrichments were identified in the posterior, with 54 early and 40 late DEGs (Fig. 2F). The magnitude of the expression changes was also modest, with the majority of DEG effect sizes (absolute value of the log2 fold change) of two or less (59% of estrous cycle DEGs, vs. 44% of AP DEGs). This result contrasts starkly to the transcriptional dynamics in the oviducts of other organisms, including human, bovine, and swine, across the menstrual/estrous cycle (Bauersachs et al., 2004; Cerny et al., 2015; Hess et al., 2013; Kim et al., 2018). For example, in bovine anterior oviducts, 972 genes have been reported as enriched early and 597 enriched late (Cerny et al., 2015), while in human anterior oviducts, 650 genes have been reported as enriched early and 683 enriched late (Hess et al., 2013).

In contrast to the muted transcriptional dynamics across the estrous cycle, we observed highly robust and spatially-restricted patterns of gene expression along the AP axis of the mouse oviduct. We identified 1337 genes enriched in the anterior, and 770 in the posterior early in the estrous cycle (Fig. 2G). Similarly, late in the estrous cycle, 1579 genes were enriched in the anterior and 975 in the posterior (Fig. 2H). Thus, our data suggest that the mouse oviduct experiences relatively modest transcriptional changes as it progresses through the estrous cycle, but displays robust transcriptional differences along the AP axis.

### The mouse oviduct transcriptome is relatively stable across the estrous cycle

While our finding of modest transcriptional variation across the estrous cycle in the mouse oviduct was consistent with the general lack of robust tissue remodeling, it was still surprising given the impact of cycling steroid hormones on oviduct function (Barton et al., 2020). To ask if the small number of DEGs we observed across time were enriched in particular classes, we performed Gene Ontology (GO) analysis (Alexa, 2020) on the early versus late DEGs. In the late stages, DEGs were subtly enriched for xenobiotic metabolism: xenobiotic metabolic process, cellular response to xenobiotic stimulus, and xenobiotic catabolic process (Supp. Fig. 1B).

More interestingly, DEGs in the early stages were enriched for translation-related terms: peptide metabolic process, peptide biosynthetic process, translation, gene expression, and cellular amide metabolic process (Supp. Fig. 1A). This enrichment for translation-related terms early in the estrous cycle is consistent with the known concomitant increase in protein concentration (Bronson and Hamilton, 1971). In agreement with the GO analysis, manual annotation revealed an enrichment of large and small ribosome subunit genes expressed in the early estrous cycle compared to late (Fig. 3A), with these genes displaying a 2 to 4-fold change in abundance (Fig. 3B).

**Figure 3.**
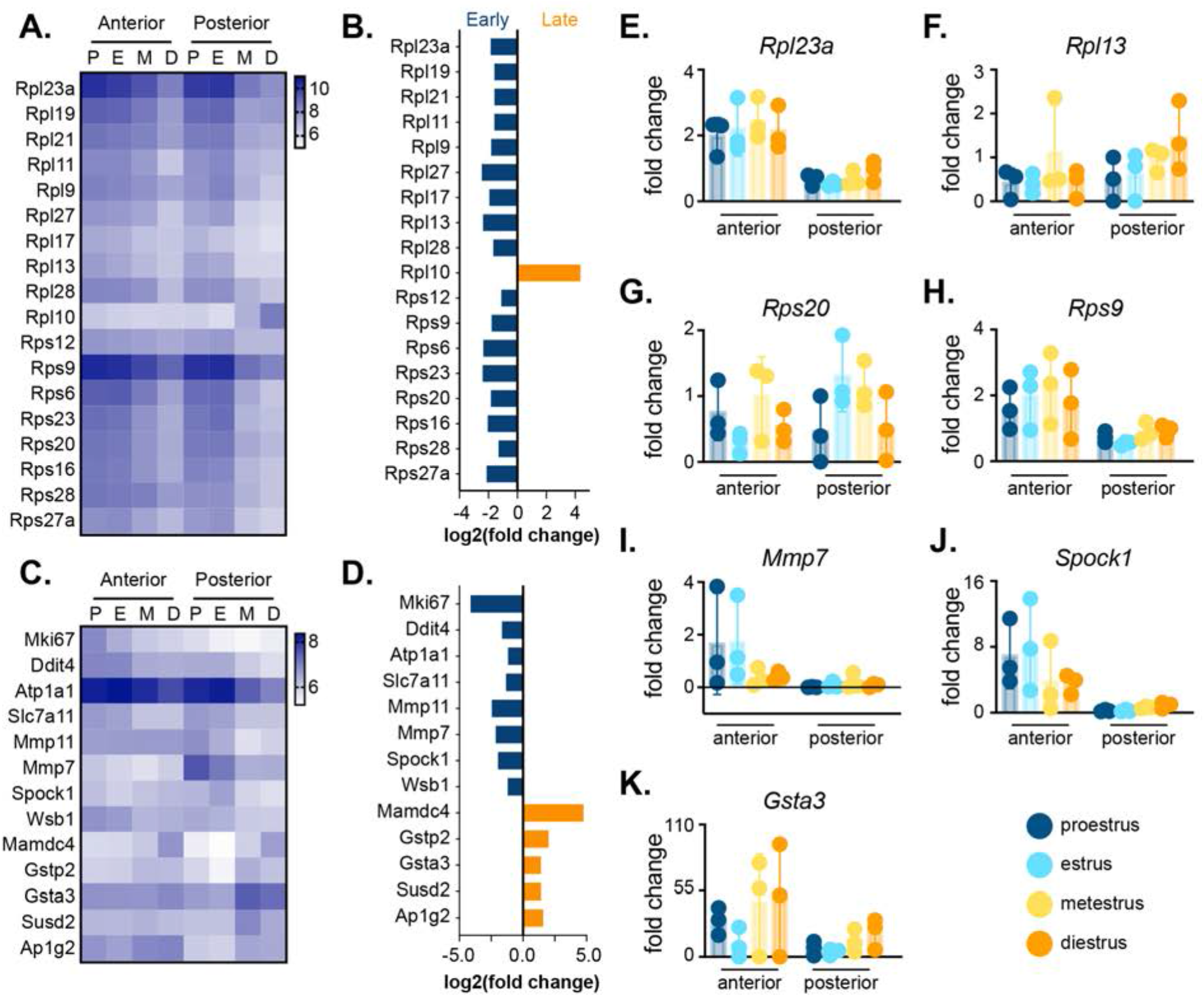
The mouse oviduct transcriptome is remarkably stable across the estrous cycle. A) Heatmap depicting expression of ribosomal genes across the estrous cycle where rows indicate genes and columns indicate the estrous cycle stage (P-proestrus, E-estrus, M-metestrus, D-diestrus). B) The effect size (absolute value of the log2 fold change) of the ribosomal genes was generally modest (under 2). C) Heatmap of other DEGs. D) These genes also generally have modest but consistent effect sizes. E-K) qPCR histograms of select genes from B and D along the anteroposterior axis at each estrous cycle stage. All heatmap scales use DESeq2 variance stabilized counts.

Further curation identified potentially interesting candidates based on their known or hypothesized function in female fertility. For example, *Mki67*, *Ddit4*, *Atp1a1*, *Slc7a11*, *Mmp11*, *Mmp7*, *Spock1*, and *Wsb1* were each enriched at early stages, while *Mamdc4*, *Gstp2*, *Gsta3*, *Susd2*, and *Ap1g2* were enriched late, either in the anterior, posterior, or both (Fig. 3C). Among these, *Mmp7* and *Slc7a11* were especially interesting, because both were upregulated early and down regulated late, a pattern that is consistent with that observed in the cycling human oviduct (Hess et al., 2013). *Mmp7* is part of the matrix-degrading enzyme family, and its down-regulation late in the human menstrual cycle is hypothesized to help maintain oviduct matrix integrity as the pre-implantation embryo travels through the organ (Hess et al., 2013). *Slc7a11* is a glutamate/cysteine antiporter that is similarly regulated in the bovine oviduct (Cerny et al., 2015). The majority of these DEGs also displayed modest effect sizes, similar to the ribosomal genes (Fig. 3D).

To independently verify the observed stability of the oviduct transcriptome across the estrous cycle, we performed qPCR for several of the DEGs. Consistent with the transcriptome-wide data, we did not observe statistically significant changes across the estrous cycle for most genes assayed (Fig. 3E-K), but the trends in expression consistently reflected trends in our TagSeq data. The robust differential expression observed by qPCR between anterior and posterior (see next section) indicate that these negative findings across the cycle do not reflect a lack of sensitivity in our assays. Rather, we conclude that the relatively modest remodeling observed in the oviduct by histology (Fig. 1) is reflected by relatively muted transcriptional dynamics across the estrous cycle. While future work will be required, it seems reasonable to suggest that the stability of the mouse oviduct across the estrous cycle represents an evolutionarily derived state associated with the extremely rapid mouse estrous cycle, which at four to five days, is much faster than in other organisms (Brenner and West, 1975).

### The mouse oviduct displays robust transcriptional patterning along the anteroposterior axis

We next sought to understand the robust changes in gene expression we observe along AP axis of the oviduct. We therefore turned again to GO Term analysis (Alexa, 2020). Consistent with the strong anterior enrichment of MCCs observed by histology (Fig. 1), anterior-enriched DEGs were highly enriched for cilia-related processes (Supp. Fig. 1C). In the anterior, complexes that function in MCCs, including intraflagellar transport, transition zone, CPLANE, and BBSome proteins (Garcia et al., 2018), as well as ciliary motility (Legendre et al., 2020) and MCC transcription factors (Lewis and Stracker, 2020) were enriched (Fig. 4A). To provide additional resolution to these patterns of expression for MCC-specific genes, we performed qPCR for several at each estrous cycle stage, using not just anterior and posterior regions but also the intervening middle region of the oviduct. We observed robust, statistically significant enrichment in the anterior region for a ciliary transcription factor (*Foxj1*), a transition zone complex gene (*Tmem231*), two ciliogenesis genes (*Ift140* and *Ift57*), and one gene encoding a component of the ciliary beating machinery (*Dnah9*) (Fig. 4F-I).

**Figure 4.**
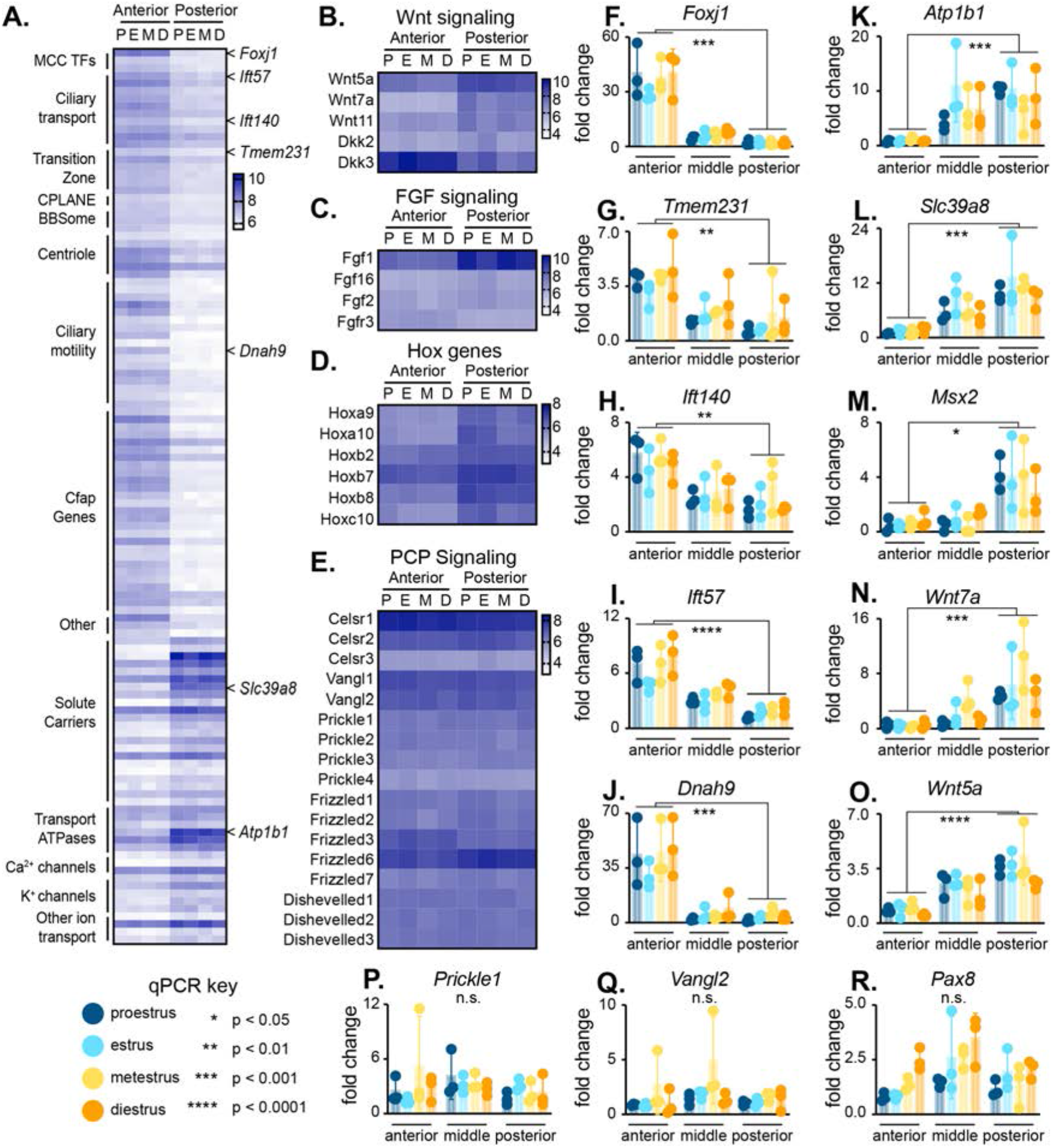
The mouse oviduct displays robust transcriptional patterning along the anteroposterior axis. A) Heatmap showing expression dynamics of DEGs across the anterior and posterior, including cilia-related genes and complexes, solute carriers, transport ATPases, and calcium and potassium channels. Additional heatmaps of genes in developmental signaling pathways, including B) Wnt signaling, C) FGF signaling, D) Hox genes, and E) PCP signaling. F-R) qPCR histograms of select genes from A-E along the anteroposterior axis at each estrous cycle stage: anterior-enriched genes *Foxj1*, *Tmem231*, *Ift140*, *Ift57*, *Dnah9;* posterior-enriched genes *Atp1b1*, *Slc29a8*, *Msx2*, *Wnt7a*, *Wnt5a*; and other qPCR assayed genes *Prickle1*, *Vangl2*, *Pax8*.

To investigate whether mouse anterior oviduct DEGs are similar to human multiciliated tissues, we compared our MCC-enriched dataset to the recently published human multiciliated tissue transcriptional signature (Patir et al., 2020). The analysis of four human multiciliated tissues, including ependymal, oviduct, trachea, and sperm, identified 248 genes expressed by all four tissues. Of those, we find 166 of them to be differentially expressed in the anterior of the mouse oviduct (data not shown), suggesting that, transcriptionally, multiciliated genes of the mouse oviduct are highly similar to those of human multiciliated tissues.

Non-ciliated epithelial cells in the posterior oviduct consist predominantly of secretory cells (Agduhr, 1927; Ghosh et al., 2017). Accordingly, our posterior-enriched gene set was strongly enriched for transport and secretion GO Terms (Supp. Fig. 4), including solute carriers, transport ATPases, Ca^2+^ and K^+^ channels (Fig. 4A). These GO terms are similar to previously published datasets from other mammals, including cows and humans, where vesicle-mediated transport, endocytosis, and exocytosis were among the main processes enriched in the posterior (Gonella-Diaza et al., 2017; Maillo et al., 2016; Rose et al., 2020). We confirmed this trend using qPCR for a transport ATPase (*Atp1b1*) and a solute carrier (*Slc39a8*); both were significantly enriched in the posterior (Fig. 4K, L).

Interestingly, certain genes enriched in the mouse oviduct posterior are differentially expressed across the human menstrual cycle (Hess et al., 2013). For example, of the 33 solute carrier (Slc) genes that are enriched in the posterior of the mouse oviduct, eight of them are also differentially expressed across the human menstrual cycle, including *Slc2a3*, *Slc22a23*, *Slc27a3*, *Slc39a14*, *Slc39a2*, *Slc39a8*, *Slc4A7*, *Slc8a1* (Hess et al., 2013). These Slc genes are hypothesized to be important for secreting amino acids and other nutrients into the oviductal lumen to aid in preimplantation embryonic development. Further studies investigating SLC expression dynamics across both humans and mice may offer insights into the evolution of mammalian reproduction and divergence underlying these secretory cell types.

### Patterned expression of known developmental signaling systems along the anteroposterior axis of the adult oviduct

Our analysis of anteroposterior gene expression patterns also revealed several previously unreported trends. First, the genes up-regulated specifically in the posterior oviduct were strongly enriched for GO Terms related to embryonic development (Supp. Fig. 1D). For example, all three non-canonical Wnt ligands, *Wnt5a*, *Wnt7a*, and *Wnt11* were enriched in the posterior region of the oviduct (Fig. 4B, N, O). Moreover, two antagonists of canonical Wnt signaling were also differentially expressed, but in a curious fashion: *Dkk2* was enriched in the posterior, while *Dkk3* was enriched in the anterior (Fig. 4B). Components of the FGF signaling pathway were also differentially expressed: *Fgf1*, *Fgf16*, and *Fgf2* were enriched in the posterior, while only the FGF receptor, *Fgfr3*, was enriched in the anterior (Fig. 4C). Finally, multiple transcription factors were differentially expressed, including Msx2 (Fig. 4M) and several Hox genes, including *Hoxa9* and *Hoxa10* (Fig. 4D), both of which are involved in female reproductive tract development (Du and Taylor, 2015). Other differentially expressed Hox genes include *Hoxb2*, *Hoxb7*, *Hoxb8*, and *Hoxc10* (Fig. 4D). We confirmed the differential expression of many of the developmental regulators using qPCR (Fig. 4M-O)

Not all signaling pathways displayed patterned expression, and an excellent example is the Planar Cell Polarity (PCP) pathway (Fig. 4E). The PCP system is essential for normal polarized ciliary beating in the oviduct, so most PCP genes are expressed in the oviduct (Koyama et al., 2019; Shi et al., 2014). Strikingly however, none displayed enrichment along the AP axis (Fig. 4E).

In contrast to intracellular effectors of PCP signaling, expression of non-canonical Wnt ligands was strongly polarized, which was of interest for multiple reasons. First, Wnt signaling is necessary to maintain stemness in human oviduct organoid cultures (Kessler et al., 2015), and secretory cells are thought to be the progenitors of MCCs in the mouse oviduct. Thus, it may be that posteriorly enriched Wnt signaling maintains the progenitor capabilities of posteriorly enriched secretory cells (Ghosh et al., 2017). Second, *Wnt5a*/*7a*/*11* are known to orient PCP signaling, providing directional cues for the apical surface of cells and driving coordinated ciliary beating in multiciliated tissues (Butler and Wallingford, 2017; Gao et al., 2011; Koyama et al., 2019; Ossipova et al., 2015). While the oviduct is planar polarized – i.e. Vangl2 localizes anteriorly in oviduct epithelial cells – it is unknown where the directional cue originates (Shi et al., 2016). Our data raise the possibility that posteriorly enriched *Wnt5a/7a/11* orients Vangl2 localization to the anterior side of the oviduct epithelium, thereby regulating ciliary flow towards the posterior oviduct. Our experiments also indicate that PCP signaling components are evenly expressed across the AP axis of the oviduct, While the impact of planar polarization on MCCs is fairly well studied, the corresponding impact on secretory cells remains unknown.

## Conclusions

In summary, our transcriptome profiling data fills a major gap in mouse oviduct investigations. While confirming reports made using other methods, our detailed anteroposterior axis and estrous stage analyses reveal novel gene expression patterns as well as providing a foundation for further studies. Hypotheses generated from these data can inform additional explorations using orthogonal methods such as single-cell sequencing and proteomics. Such studies have the potential to identify specific cell types associated with expression trends observed here, as well as quantifying actual cellular protein abundances which may differ from RNA expression.

## Acknowledgements

We thank the Genome Sequencing and Analysis facility at the University of Texas at Austin for their support and assistance with sequencing. This research was supported by funding from the National Institutes of Health (F32HD095618 to E.C.R., R01HL117164 to J.B.W., R01HD085901 to J.B.W. and E.M.M., and R35GM122480 to E.M.M.), the American Heart Association predoctoral fellowship (#18PRE34060258) to R.K.G., and the Welch Foundation (F-1515, to E.M.M.).

---

**Supplementary Figure 1.**
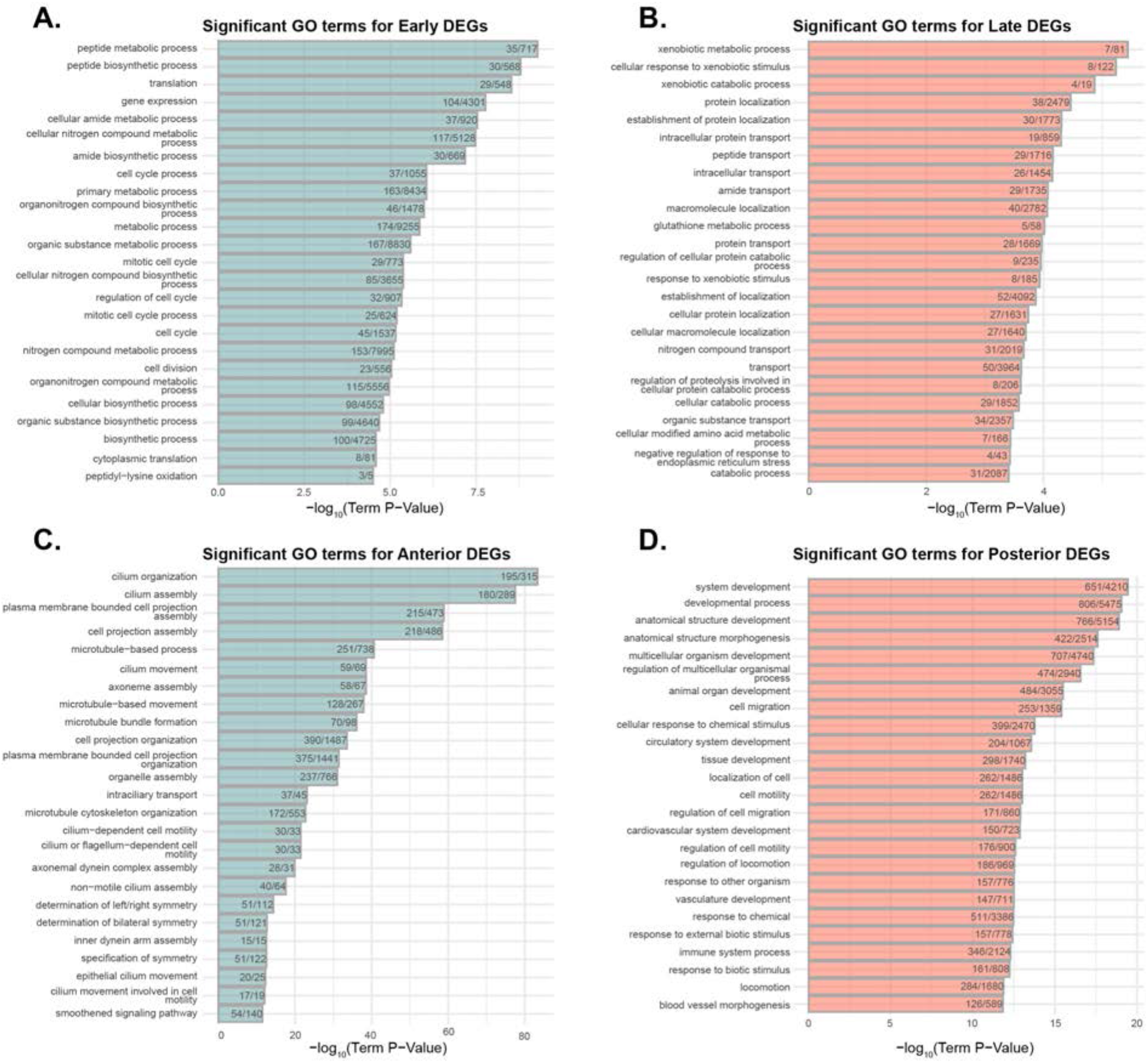
Significant GO terms associated with oviduct transcriptome analysis. Histograms depicting significant GO terms for A) early, B) late, C) anterior, and D) posterior DEGs.

## Supplemental spreadsheet tabs

**S-1. Primers used for qPCR.**

**S-2. Description of the sample datasets used.**

Full DEG and GO term details are available in the Supplementary zip file for GEO accession number GSE164718.

---

**Table S-1.**
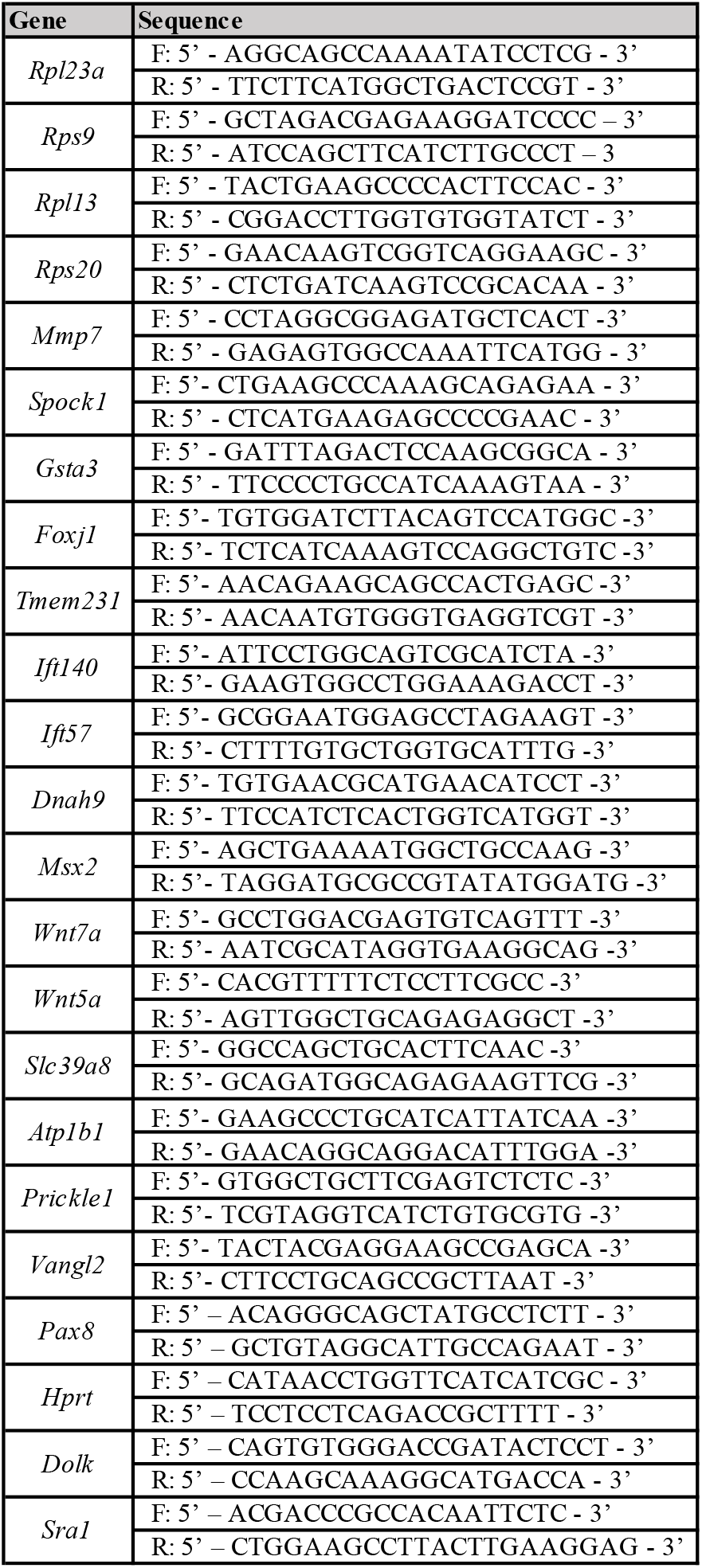

**Table S-2.**

| Table S-2 |  |  |  |  |  |  |  |  |  |  |  |
| --- | --- | --- | --- | --- | --- | --- | --- | --- | --- | --- | --- |
| ID | Sample | Location | Stage | Phase | Location Phase | # Raw reads | # Deduped | # Aligned | % Align | GEO Fastq file | GEO kallisto counts file |
| S1 | Proes Ant B1 | Anterior | Proestrus | Early | Anterior Early | 2,669,590 | 1,392,774 | 1,038,851 | 74.6% | Proes Ant B1 R1.fastq.gz | Proes Ant B1 kallisto counts.tsv |
| S2 | Proes Ant B2 | Anterior | Proestrus | Early | Anterior Early | 2,903,086 | 1,583,735 | 1,184,664 | 74.8% | Proes Ant B2 R1.fastq.gz | Proes Ant B2 kallisto counts.tsv |
| S3 | Proes Post B1 | Posterior | Proestrus | Early | Posterior Early | 3,820,771 | 1,809,899 | 1,358,747 | 75.1% | Proes Post B1 R1.fastq.gz | Proes Post B1 kallisto counts.tsv |
| S4 | Proes Post B2 | Posterior | Proestrus | Early | Posterior Early | 2,542,319 | 1,258,262 | 936,809 | 74.5% | Proes Post B2 R1.fastq.gz | Proes Post B2 kallisto counts.tsv |
| S5 | Es Ant B1 | Anterior | Estrus | Early | Anterior Early | 3,859,515 | 2,015,223 | 1,524,157 | 75.6% | Es Ant B1 R1.fastq.gz | Es Ant B1 kallisto counts.tsv |
| S6 | Es Ant B2 | Anterior | Estrus | Early | Anterior Early | 4,976,661 | 2,393,539 | 1,803,496 | 75.3% | Es Ant B2 R1.fastq.gz | Es Ant B2 kallisto counts.tsv |
| S7 | Es Post B1 | Posterior | Estrus | Early | Posterior Early | 3,051,253 | 1,508,916 | 1,166,422 | 77.3% | Es Post B1 R1.fastq.gz | Es Post B1 kallisto counts.tsv |
| S8 | Es Post B2 | Posterior | Estrus | Early | Posterior Early | 1,486,318 | 792,559 | 589,353 | 74.4% | Es Post B2 R1.fastq.gz | Es Post B2 kallisto counts.tsv |
| S9 | Metes Ant B1 | Anterior | Metestrus | Late | Anterior Late | 1,509,395 | 896,284 | 696,932 | 77.8% | Metes Ant B1 R1.fastq.gz | Metes Ant B1 kallisto counts.tsv |
| S10 | Metes Ant B2 | Anterior | Metestrus | Late | Anterior Late | 1,742,088 | 1,000,503 | 771,609 | 77.1% | Metes Ant B2 R1.fastq.gz | Metes Ant B2 kallisto counts.tsv |
| S11 | Metes Post B1 | Posterior | Metestrus | Late | Posterior Late | 3,034,943 | 1,266,394 | 989,728 | 78.2% | Metes Post B1 R1.fastq.gz | Metes Post B1 kallisto counts.tsv |
| S12 | Metes Post B2 | Posterior | Metestrus | Late | Posterior Late | 4,033,746 | 1,918,652 | 1,484,995 | 77.4% | Metes Post B2 R1.fastq.gz | Metes Post B2 kallisto counts.tsv |
| S13 | Dies Ant B1 | Anterior | Diestrus | Late | Anterior Late | 4,403,124 | 2,067,053 | 1,605,618 | 77.7% | Dies Ant B1 R1.fastq.gz | Dies Ant B1 kallisto counts.tsv |
| S14 | Dies Ant B2 | Anterior | Diestrus | Late | Anterior Late | 4,161,010 | 1,439,043 | 1,088,609 | 75.6% | Dies Ant B2 R1.fastq.gz | Dies Ant B2 kallisto counts.tsv |
| S15 | Dies Post B1 | Posterior | Diestrus | Late | Posterior Late | 2,131,914 | 946,725 | 726,724 | 76.8% | Dies Post B1 R1.fastq.gz | Dies Post B1 kallisto counts.tsv |
| S16 | Dies Post B2 | Posterior | Diestrus | Late | Posterior Late | 1,522,405 | 721,007 | 572,923 | 79.5% | Dies Post B2 R1.fastq.gz | Dies Post B2 kallisto counts.tsv |

## Notes

### Competing Interest Statement

The authors have declared no competing interest.

### Summary of Updates

This revision corrects a mis-spelling of Riddhiman Garge's name in the Author list.

